# KLF7-Regulated ITGA2 as a Therapeutic Target for Inhibiting Oral Cancer Stem Cells

**DOI:** 10.1101/2024.11.04.621805

**Authors:** Xin Qi, Jiang Zhou, Pan Wang, Yunyan Li, Haoran Li, Yuwen Miao, XiaoQing Ma, Xiayan Luo, Zhiling Zhang, Yanling He, Wenyi shen, Wenquan Zhao, Rutao Cui, Cang Li, Huiyong Zhu, Jiong Lyu

**Author notes:** These authors contributed equally: Xin Qi, Jiang Zhou, Pan Wang. These authors jointly supervised this work: Cang Li, Huiyong Zhu, and Jiong Lyu. e-mail.

## Abstract

Cancer stem cells (CSCs) play crucial roles in tumor metastasis, therapy resistance, and immune evasion. Identifying and understanding the factors that regulate the stemness of tumor cells presents promising opportunities for developing effective therapeutic strategies. In this study on oral squamous cell carcinoma (OSCC), we confirmed the key role of KLF7 in maintaining the stemness of OSCC. Using chromatin immunoprecipitation sequencing and dual-luciferase assays, we identified ITGA2, a membrane receptor, as a key downstream gene regulated by KLF7 in the maintenance of stemness. Tumor sphere formation assays, flow cytometry analyses, and in vivo limiting dilution tumorigenicity evaluations demonstrated that knocking down ITGA2 significantly impaired stemness. When bound to its ECM ligand, type I collagen, ITGA2 activates several stemness-related pathways, including PI3K-AKT, MAPK, and Hippo. TC-I 15, which inhibits the ITGA2–collagen interaction, showed a synergistic anti-tumor effect when combined with cisplatin in both *in vitro* and xenograft models. In summary, we reveal that the KLF7/ITGA2 axis is a crucial modulator of stemness in OSCC. Our findings suggest that ITGA2 is a promising therapeutic target, offering a novel anti-CSC strategy.

**Highlights:**

1) KLF7 as a key molecule in maintaining oral cancer stemness.
2) ITGA2as a key downstream gene regulated by KLF7 in the maintenance of stemness.
3) ITGA2 interacts with extracellular matrix type I collagen, activating stemness-related pathways and promoting YAP1 nuclear translocation to sustain OCSCs.
4) ITGA2 as a novel anti-CSC target, providing a new strategy to overcome OSCC drug resistance.

## Introduction

Oral cancer affects the buccal mucosa, floor of the mouth, oral tongue, alveolar ridge, retromolar trigone, and hard palate. It has an incidence rate of approximately 1 in 100,000 individuals^1^. The most common pathological type is oral squamous cell carcinoma (OSCC), accounting for more than 90% of cases^2^. Early-stage OSCC (Stages I–II) is typically treated with surgery alone, whereas advanced OSCC (Stages III-IV) requires multimodal treatment approaches and has a relatively poor prognosis due to its tendency for recurrence and lymph node metastasis^3^. Due to low early diagnosis rates, most OSCC patients are already at an advanced stage at the time of diagnosis^4^. Current treatments for advanced OSCC primarily include surgery, chemotherapy, radiotherapy, or a combination of these approaches. Despite the various treatments employed over the decades, the overall survival rate for OSCC remains at 50%^5^. Common chemotherapy agents for OSCC include platinum drugs, 5-fluorouracil (5-FU), paclitaxel (PTX), and doxorubicin (Dox). However, the majority of patients develop resistance to these drugs. At present, multidrug resistance is one of the primary obstacles to successful cancer chemotherapy and leads to poor patient prognosis^6^.

The concept of cancer stem cells (CSCs) has evolved over several decades^7^. They were first identified in the 1990s in hematologic malignancies and have since been isolated from various hematologic and solid tumors, including oral cancer^8–12,13^. Oral cancer stem cells (OCSCs) have the capacity for long-term self-renewal and can replicate the diverse cell lineages found in primary cancers^14,15^. One characteristic of OCSCs is their propensity to develop resistance to treatments, including both conventional chemotherapy and immunotherapy. Therefore, therapies targeting OCSCs hold promise as an effective approach to overcoming resistance in OSCC. Developmental signaling pathways, including Hippo, Notch, and WNT, are frequently altered in CSCs and play key regulatory roles in supporting stem cell maintenance and survival^16^. Various CSC-targeted therapies have been developed and are currently in clinical trials, such as NEDD8-activating enzyme inhibitors targeting the Hippo pathway, γ-secretase inhibitors targeting the Notch pathway, and small-molecule antagonists of β-catenin targeting the WNT pathway^17–21^. Progress has also been made in targeting OCSCs in oral cancer. For example, Li et al. demonstrated that β-catenin silencing enhances OSCC sensitivity to cisplatin^22^. Additionally, CD133 expression is elevated in OSCC and associated with increased resistance; targeting CD133 combined with cisplatin treatment effectively inhibits OCSC-driven OSCC initiation^23^.

Utilizing novel drugs to target therapy-resistant CSCs may reduce cancer recurrence rates and improve treatment efficacy. To mitigate drug resistance in OSCC, it is essential to improve knowledge of OCSCs, with a particular focus on molecular features. Recent evidence suggests that CSCs represent a plastic cellular state influenced by dynamic interactions within the CSC niche, rather than a fixed condition^24^. Differentiated cancer cells can revert to a more undifferentiated state under certain conditions or stimuli^25^. The maintenance of CSCs depends on the tumor microenvironment and niche. In this study, we employed single-cell analysis to identify a stem-like subset within OSCC, characterized by high developmental potential and activation of stemness pathways. We identified KLF7 as a potential key molecule in maintaining OCSC. Further investigation revealed that KLF7 regulates OSCC stemness through the direct transcriptional activation of ITGA2, a membrane protein crucial for mediating cell-ECM interactions. ITGA2 inhibition can suppress OSCC stemness and exhibits a synergistic anti-tumor effect when combined with cisplatin. Our findings highlight that ITGA2 is a promising therapeutic target, offering a novel anti-CSC strategy.

## Results

### Identification of CSC and key molecules for maintaining stemness in OSCC

To identify the key molecule of CSCs in OSCC, we analyzed single-cell transcriptome data from 10 OSCC samples obtained from public databases (Supplementary Fig.2a). After a series of preprocessing steps, we annotated 55,000 cells using previously published marker gene sets^26–28^ (Fig.1a and Supplementary Fig.1b). Using HoneyBadger to exhibiting high copy-number variation (Supplementary Fig.1c). Epithelial-derived cells with CNV >0.1 were identified as malignant cells and further subdivided into subtypes. Unsupervised Louvain-based clustering and UMAP identified four distinct OSCC cell states and GSVA functionally annotated each cluster, and S1 was identified as the stem-like subtype (Fig.1b,c and Supplementary Tab. 1). S1 exhibited the highest developmental potential, positioned at the beginning of the cell trajectory, and showed the highest stemness gene expression activity(ref^29^)(Fig.1d,e and Supplementary Fig.1d). Consistent with studies linking CSCs and tumor metastasis^30^, we found that S1 exhibited the highest invasive characteristics(ref^31^)(Supplementary Fig.1e). These findings indicate that the cells within S1 possess the highest stem-like properties among all tumor cell.

**Fig. 1:**
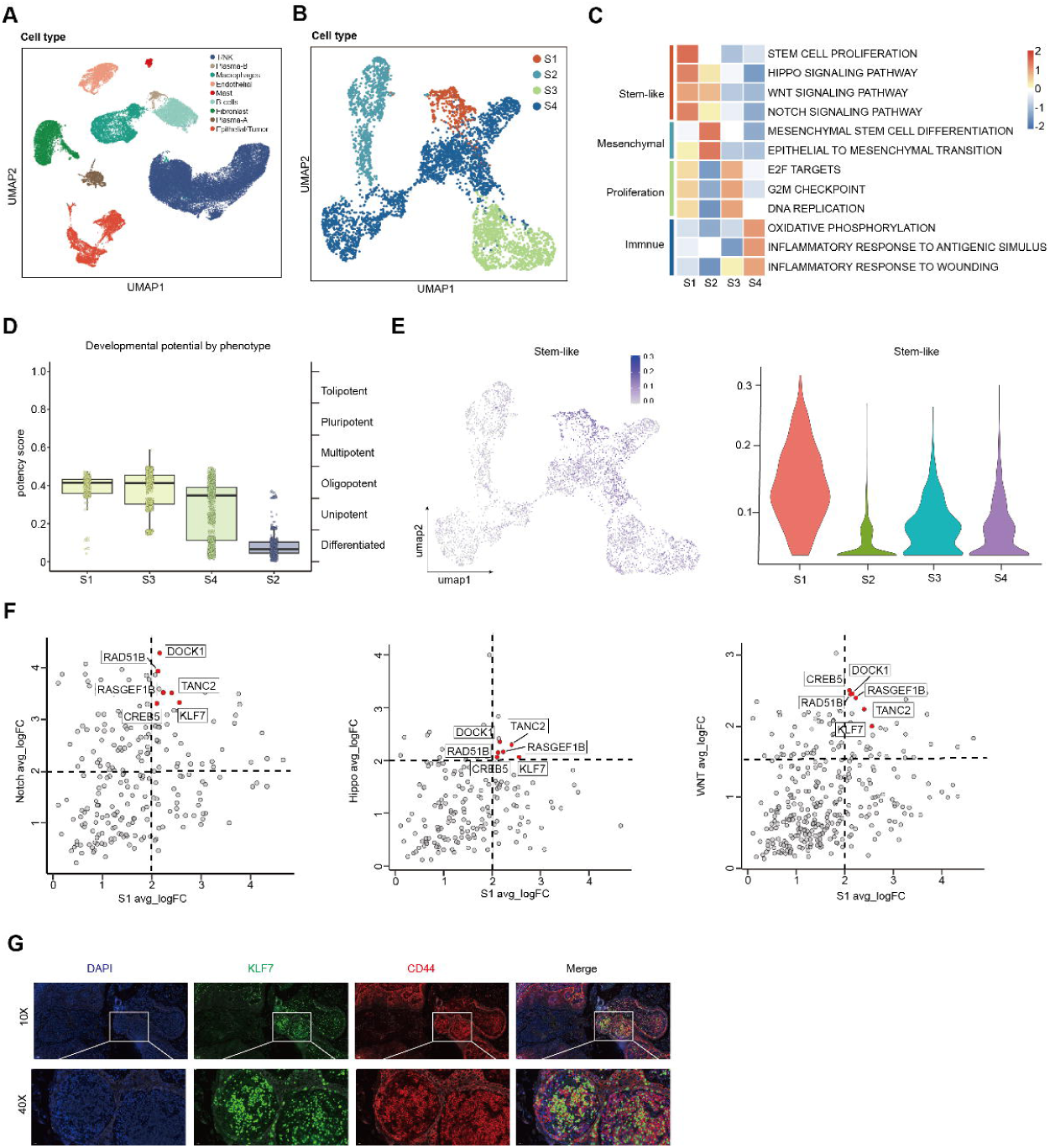
Identification of Cancer Stem Cells and Key Molecules for Maintaining Stemness in Oral Cancer. **A** UMAP illustrating cell type diversity in human OSCC(50k cells). **B** UMAP plot of the subtype of malignant cells analysed by scRNA-seq. **C** Heatmap depicting enrich pathway of the subtype of malignant cells. **D** Cytorace plot of the subtype of malignant cells. **E** Stem-like gene sets activity (AUCell score) in all malignant cells (umap plots, left panel) or per transcriptional state (violin plots, right panel). **F** The log2-transformed fold change (FC) in RNA levels between the stem-like cluster with the rest of the clusters and the high enrichment in the stem-related pathway. **G** Representative immunofluorescent image showing the expression of KLF7, CD44, and DAPI in human OSCC tissue.

We sought to identify the key molecule in the stem-like subtype. It is well-established that maintaining stemness requires the activation of multiple signaling pathways. We found six molecules highly enriched in the stem-like subtype and stem-related pathway (Notch, Wnt, and Hippo pathways) (Fig.1f and Supplementary Tab.1). Notably, KLF7 is highly expressed across various tumors, with particularly high expression in head and neck squamous cell carcinoma (Supplementary Fig. 2a). It is overexpressed in OSCC (Supplementary Fig. 2b, c) and is linked to poor prognosis (Supplementary Fig. 2d). Recent research has shown that KLF7 can re-establish the hematopoietic stem cell niche, suggesting that KLF7 may be a key molecule for maintaining stemness in OSCC^32^. Furthermore, Immunofluorescence staining of OSCC samples revealed a consistent distribution of KLF7 and CD44^33,34^(Fig.1g). These findings suggest that elevated KLF7 expression is linked to the stemness properties of OSCC.

#### KLF7 regulates OSCC stemness and metastasis

To further confirm whether KLF7 regulates stemness in OSCC, we established KLF7-knockdown OSCC cell lines (CAL27 and HSC3)(Fig.2a). In vivo limiting dilution tumorigenicity assays using CAL27 cells, we observed that KLF7 knockdown significantly impaired the tumorigenicity compared to controls (Fig.2b). KLF7 knockdown resulted in reduced NANOG expression, smaller tumor spheres, and a decreased proportion of CD133+(CSC marker) cells in both CAL27 and HSC3 cell lines(Fig.2a-f)^35^.

**Fig. 2:**
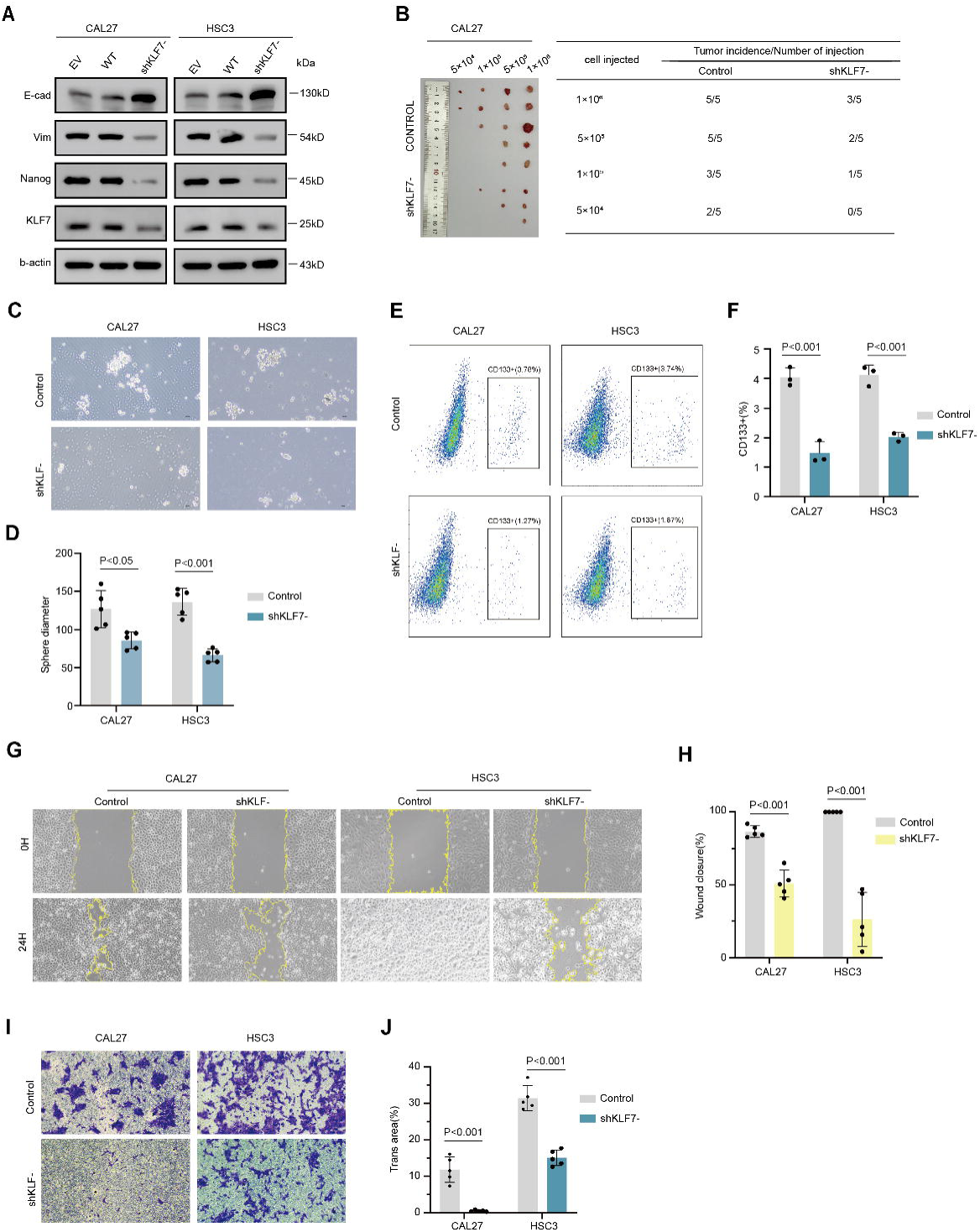
Knockdown KLF7 inhibits the stemness and metastasis of OSCC. **A** Immunoblot assay of KLF7, NANOG, VIM, and e-cad protein levels in CAL27 and HSC3 cells after stable silencing KLF7. **B** In vivo limiting dilution tumorigenicity assays of wt and silencing KLF7 CAL27 cells. The frequency of allograft formation at each injected cell dose is shown. **C** Representative image of Sphere formation in wt and silencing KLF7 cells in normal sphere medium (*n =* 5). **D** Quantification of spheres diameter in C. **E** Identification of OSCC CSCs (CD133+) in wt and silencing KLF7 cells by flow cytometry (*n =* 3). **F** Quantification of FACS analysis in E**. G** Representative image of wound-healing image after 24h in wt and silencing KLF7 cells (n=5). **H** Quantification of the percentage of wound closure in G. **I** Representative image of transwell after 24h in wt and silencing KLF7 cells (n=5). Representative image of transwell after 24h. **J** Quantify the percentage of trans area in I.

Given the reported overlap between epithelial-mesenchymal transition (EMT) and CSC characteristics^36^, we analyzed the relationships between KLF7, migration, and EMT. KLF7 knockdown was associated with reduced wound healing rates, fewer migrating cells, decreased vimentin expression, and increased E-cadherin expression (Fig.2a,g-j). Conversely, KLF7-overexpressing CAL27 and HSC3 cell lines exhibited the opposite effects: KLF7 overexpression resulted in increased NANOG expression, a higher proportion of CD133+ cells, larger tumor spheres, and enhanced EMT traits (Supplementary Fig.3a-i). Our results suggest that KLF7 plays a crucial role in regulating both OSCC stemness and EMT.

#### The stemness-related gene ITGA2 is downstream of KLF7

KLF7 is a transcription factor capable of activating downstream genes, but it poses challenges for direct drug targeting^37^. To identify downstream target genes with therapeutic potential, we conducted chromatin immunoprecipitation sequencing (ChIP-seq) analysis, focusing on proteins that may be suitable for drug development. After processing the data, we identified 1,900 peaks with q-values < 0.05 (Supplementary Tab.2). Next, we screened for peaks located in the promoter regions of protein-coding genes (Fig.3a). A total of 179 genes were enriched in the stem-like subset and stemness-related pathways, with six genes identified in ChIP-seq showing peaks located in promoter regions (Fig.3b). Notably, ITGA2 was specifically enriched in the stem-like subset (Fig.3c). Furthermore, ITGA2 is highly expressed in OSCC and is associated with poor prognosis (Fig.3d-f). Our results suggest that ITGA2 may be the downstream gene to maintain stemness in OSCC.

**Fig. 3:**
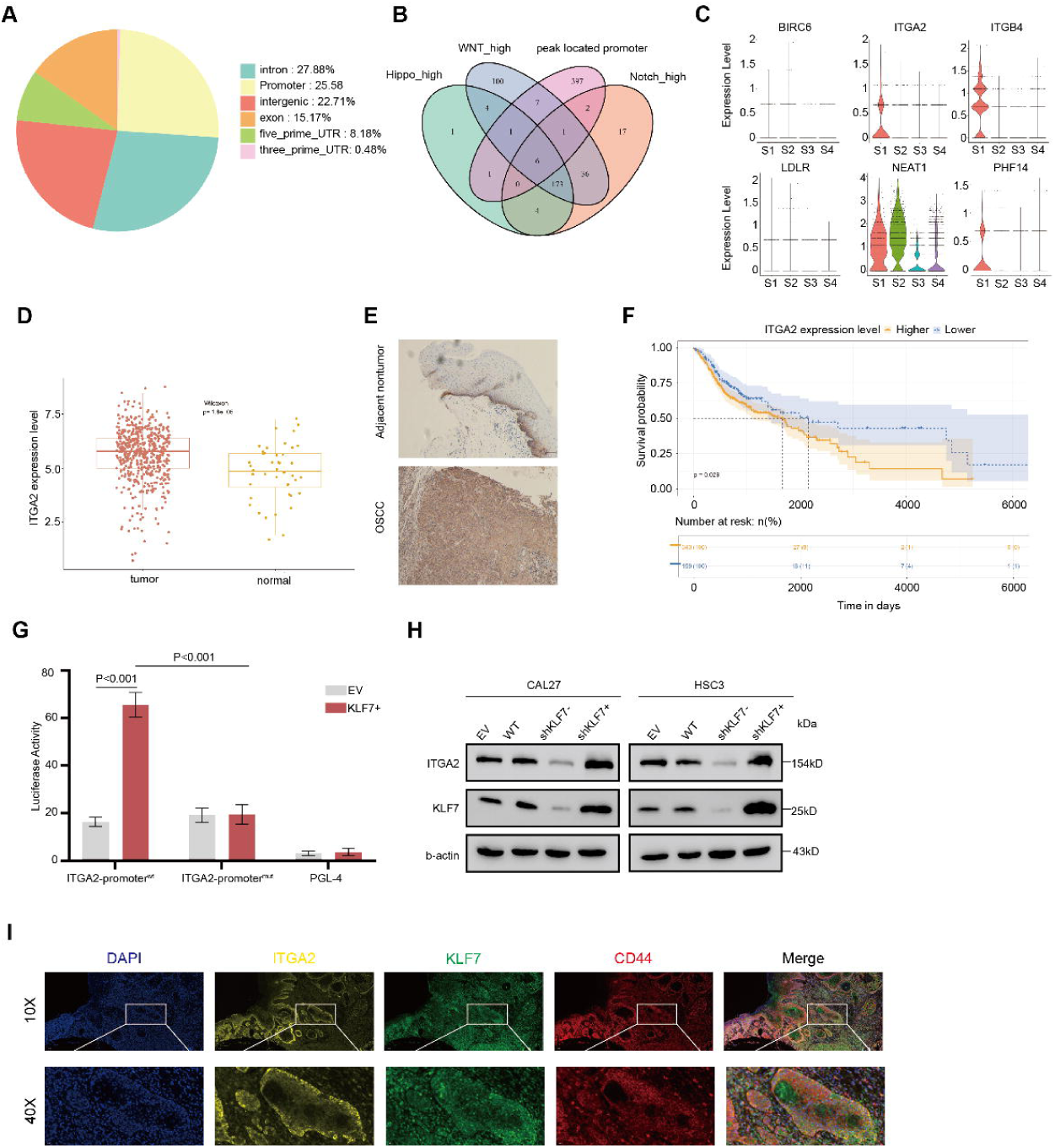
The stem-related ITGA2 is downstream gene of KLF7. **A** The distribution of BATF ChIP-seq peaks. **B** The overlap genes of highly enriched in stem-like subtype, stem-related pathway, and peaks located in the promoter region. **C** Vlnplot of overlap genes expression in OSCC subtype. **D** ITGA2 expression in OSCC and normal samples. **E** Representative immunohistochemistry image showing the expression of ITGA2 in human OSCC sample. **F** Kaplan-Meier curve of ITGA2 in OSCC TCGA dataset. **G** Luciferase activities of pGP4.19-ITGA2 promoter, mutant constructs, pRL-TK vector, when co-transfected with KLF7+ vector or control. **H** Immunoblot assay of KLF7 and ITGA2 protein levels in CAL27 and HSC3 cells after stable knockdown and overexpressing KLF7. **I** Representative immunofluorescent image showing the distribution of KLF7, ITGA2, CD44, and DAPI in human OSCC tissue.

To further confirm the direct regulation of ITGA2 by KLF7, we performed luciferase reporter assays to assess transcriptional activity. Based on the peak location in the ChIP-seq data and predictions from the JSAPAR database, we identified the KLF7 binding site at the −120/−112 region of the ITGA2 promoter. We found that luciferase activity of the pGL4.19 vector, which contains ITGA2 promoter fragments, was significantly increased following transfection of HEK293T cells with the pCMV-KLF7+ vector. Conversely, no such transactivation was observed when cells were transfected with a mutant ITGA2 promoter vector(Fig.3g). Western blot analysis confirmed that ITGA2 expression is regulated by KLF7, with ITGA2 levels decreasing upon KLF7 knockdown and increasing with KLF7 overexpression (Fig.3h). Immunofluorescence staining of OSCC samples revealed a consistent distribution of ITGA2, KLF7, and CD44 further supporting the link between KLF7, ITGA2, and stemness (Fig.3i). This funding suggest that stem-related gene ITGA2 is downstream of KLF7.

#### ITGA2 regulates OSCC stemness and metastasis

To investigate whether ITGA2 regulates OSCC stemness, we established ITGA2-knockdown CAL27 and HSC3 cell lines (Fig.4a). In vivo limiting dilution tumorigenicity assays, tumorsphere formation assays, flow cytometry, wound-healing, and transwell assays demonstrated that ITGA2 knockdown resulted in reduced tumorigenicity, fewer CD133+ cells, smaller tumorspheres, slower wound closure, and decreased cell migration (Fig.4b-f and Supplementary Fig.4a-d).

**Fig. 4:**
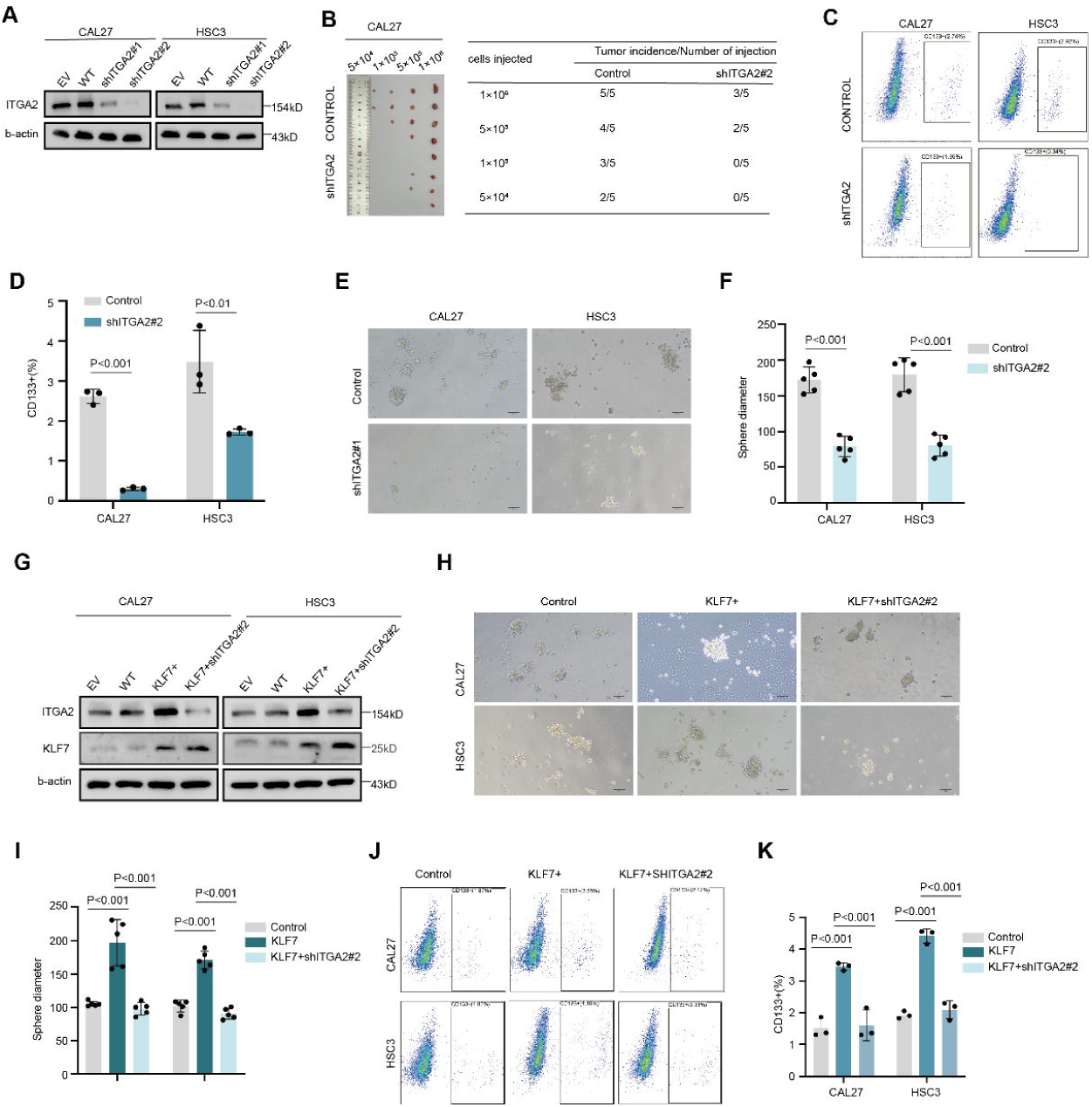
Knockdown ITGA2 inhibits the stemness of OSCC. **A** Immunoblot assay of ITGA2 protein levels in CAL27 and HSC3 cells after stable silencing ITGA2. **B** In vivo limiting dilution tumorigenicity assays of wt and silencing ITGA2 in CAL27 cells. The frequency of allograft formation at each injected cell dose is shown. **C** Identification of OSCC CSCs (CD133+) in wt and silencing ITGA2 cells by flow cytometry (*n =* 3). **D** Quantification of FACS analysis in C. **E** Representative image of Sphere formation assay in wt and silencing ITGA2 cells in normal sphere medium (*n =* 5). **F** Quantification of spheres diameter in E. **G** Immunoblot assay of KLF7 and ITGA2 protein levels after stable overexpress KLF7 and overexpress KLF7 followed by knockdown ITGA2. **H** Representative image of Sphere formation in wt, overexpress KLF7 and overexpress KLF7 followed by knockdown ITGA2 cells in normal sphere medium (*n =* 5). **I** Quantification of spheres diameter in H. **J** Identification of OSCC CSCs (CD133+) in wt, overexpress KLF7 and overexpress KLF7 followed by knockdown ITGA2 cells by flow cytometry (*n =* 3). **K** Quantification of FACS analysis in J.

Then, we knocked down ITGA2 in KLF7-overexpressing OSCC cell lines. ITGA2 knockdown attenuated the changes in tumor sphere size, CD133+ cell proportion, wound healing, and migration ability that were induced by KLF7 overexpression (Fig.4g-k and Supplementary Fig.4e-h). These findings demonstrate that the KLF7/ITGA2 axis plays a crucial role in regulating stemness in OSCC.

#### ITGA2 binds to type I collagen to drive YAP nuclear translocation and activate the stemness pathway

Although we have established that ITGA2 regulates OSCC stemness, the precise mechanisms remain unclear. In addition to embryonic pathways, PI3K-AKT and MAPK pathways also play a significant role in promoting tumor stemness. Single-cell analysis revealed that ITGA2 is enriched in the stem-like subset and is closely associated with the Hippo, MAPK, and PI3K-AKT pathways (Fig. 5a). Knockdown of ITGA2 decreased phosphorylation levels of AKT and ERK1/2, increased YAP1 phosphorylation, and led to YAP1 nuclear export(Fig.5b,c). ITGA2 contains an additional domain (“A” or “I”) within the head region, with a conserved metal ion-dependent adhesion site (MIDAS) motif which is crucial for ligand binding at the C-terminal of the type I domain(Fig.5d)^38–40^. Mutagenesis studies show that the MIDAS motif and exposed side chains on the surrounding surface are required for ligand binding^38,41–43^.

**Fig. 5:**
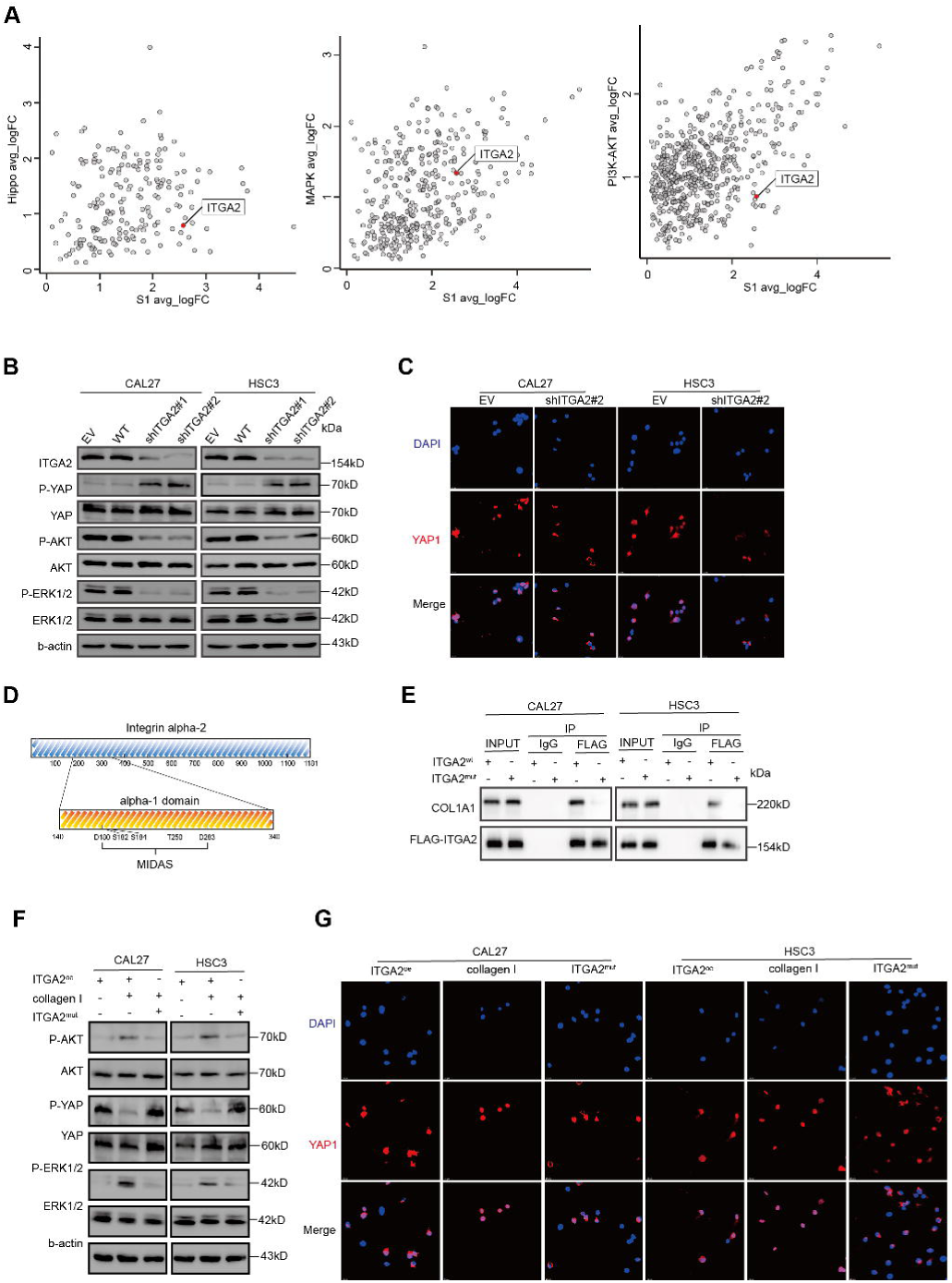
ITGA2 binds to type I collagen to drive YAP nuclear translocation and activate the stemness pathway. **A** The log2-transformed fold change (FC) in RNA levels between the stem-like cluster with the rest of the clusters and the stem-related pathway with the rest of the pathway. **B** Immunoblot assay of ITGA2, phosphor-ERK1/2, total-ERK1/2, phosphor-AKT, total-AKT, phosphor-YAP1, total-YAP1 protein levels in CAL27 and HSC3 cells after stable silencing ITGA2. **C** Representative immunofluorescent image showing the expression of YAP1 and DAPI in wt and stable silencing ITGA2 cells. **D** The sequence and domain in ITGA2 protein. **E** Co-IP analysis in CAL27 and HSC3 cells transduced with ITGA2^OE^ and ITGA2^mut^ plasmid. **F** Immunoblot assay of phosphor-ERK1/2, total-ERK1/2, phosphor-AKT, total-AKT, phosphor-YAP1, total-YAP1 protein levels in CAL27 and HSC3 cells after transduced with ITGA2^OE^ and ITGA2^mut^ plasmid. **G** Representative immunofluorescent image showing the expression of YAP1 and DAPI in CAL27 and HSC3 cells after transduced with ITGA2^OE^ and ITGA2^mut^ plasmid.

ITGA2 mediates interactions between cells and the extracellular matrix (ECM), with type I collagen as the primary ECM component^44^. To investigate whether ITGA2 interacts with type I collagen and activates stemness-related pathways, we constructed and transfected OSCC cells with ITGA2^oe^ or a ITGA2^mut^(MIDAS mutant). IP results confirmed an interaction between ITGA2 and COL1A1, which was abolished in ITGA2-mutant(Fig.5e). Upon adding type I collagen to the culture, phosphorylation of AKT and ERK1/2 increased, YAP1 decreased, and YAP1 nuclear translocation. while phosphorylation of AKT and ERK1/2 decreased, YAP1 increased and YAP1 nuclear export in ITGA2-mutant(Fig.5f,g). These findings suggest that the interaction between ITGA2 and type I collagen activates the PI3K/AKT, MAPK, and HIPPO/YAP1 pathways.

#### TC-I 15 inhibit OSCC and augments chemotherapy responsiveness of OSCC

TC-I 15 is an allosteric collagen-binding integrin α2β1 inhibitor, with half-maximal inhibitory concentration (IC50) values of 26.8 μM and 0.4 μM for binding to GFOGER and GLOGEN, respectively(Fig.6a). We selected TC-I 15 for both *in vivo* and *in vitro* experiments to evaluate the feasibility of targeting ITGA2 for OSCC treatment. Immunoprecipitation assays showed that TC-I 15 interferes with the binding between ITGA2 and type I collagen (Fig.6b). In an *in vivo* experiment, CAL27 cells were xenografted into nude mice, and from day 6 post-inoculation, TC-I 15 were intravenously injected every 3 days, with tumor volume measured over time(Fig.6c). The results showed that the TC-I 15 significantly reduced tumor size, growth rate, and tumor weight (Fig.6d). Moreover, TC-I 15 decreased phosphorylation levels of AKT and ERK1/2, increased YAP1 phosphorylation, and decreased CD133+ cells determined by flow cytometry (Fig.6e,f).

**Fig 6:**
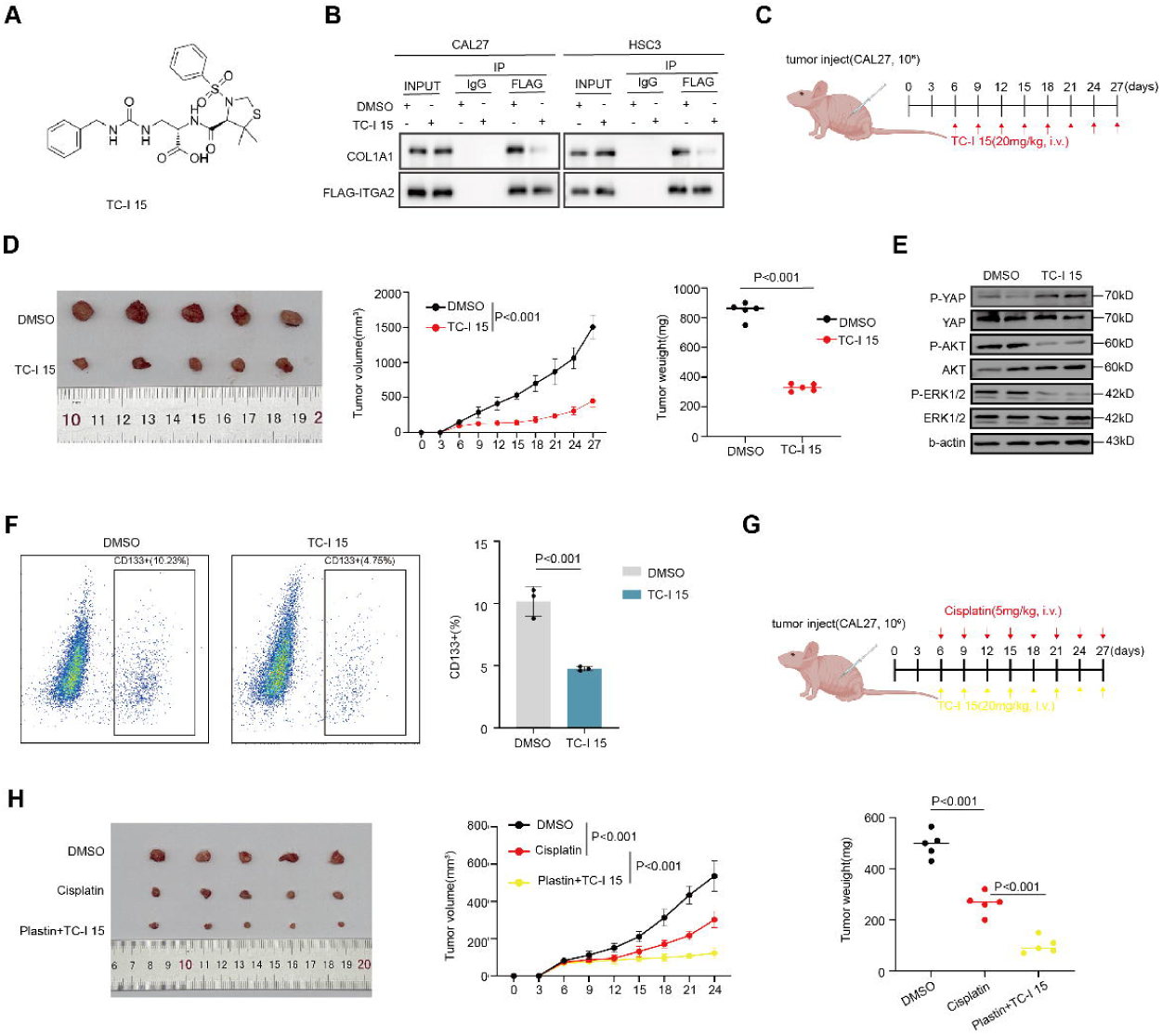
TC-I 15 inhibits OSCC and augments chemotherapy responsiveness of OSCC. **A** The molecular formula of TC-I 15. **B** Co-IP analysis in CAL27 and HSC3 cells treated with DMSO and TC-I 15. **C** CAL27 cells were intracardially injected into mice. Seven days later, TC-I 15 was injected (20 mg/kg) into mice via vein every third day for 1 month. **D** Representative images showing the xenograft model in CA27 cells treated with DMSO or TC-I 15(n = 5 per group) and the growth of tumor grafts was shown. **E** Immunoblot assay of phosphor-ERK1/2, total-ERK1/2, phosphor-AKT, total-AKT, phosphor-YAP1, and total-YAP1 protein levels in xenograft tissue. **F** Representative FACS plots and quantification of CD133+cells in the xenograft tissues. **G** CAL27 cells were intracardially injected into mice. Seven days later, TC-I 15 (20 mg/kg) and plastin(5 mg/kg) were injected into nude mice via vein every third day for 1 month. **H** Representative images showing the xenograft model in CA27 cells treated with DMSO or TC-I 15(n = 5 per group) and the growth of tumor grafts were shown.

Given the critical role of CSCs in tumor drug resistance, we further investigated the potential synergistic effects of TC-15 plus cisplatin in the treatment of OSCC. CAL27 cells were xenografted into nude mice, and from day 6 post-inoculation, cisplatin and TC-I 15 were intravenously injected every 3 days, with tumor volume measured over time(Fig.6g). The results showed that the combination of TC-I 15 and cisplatin significantly reduced tumor size, growth rate, and tumor weight compared to cisplatin alone (Fig.6h). These findings suggest that TC-I 15 inhibits OSCC and enhances the sensitivity of OSCC to chemotherapy, highlighting the potential of targeting ITGA2 to improve clinical anti-tumor efficacy.

## Discussion

Our study revealed that KLF7 modulates stemness properties in OSCC. The KLF family is closely associated with CSCs. Notably, KLF4 plays a crucial role in maintaining CSC characteristics in liver and breast cancers^45,46^. Additionally, KLF11 has been identified as a negative regulator in sarcoma CSCs, where its epigenetic silencing leads to prolonged YAP activation and poor prognosis^47^. KLF7 can also enhance cancer cell proliferation, migration, and invasion across various cancer types^48,49^. Although direct evidence linking KLF7 to CSC regulation is lacking, its ability to modulate genes involved in the EMT suggests a potential impact on CSC properties^50^. Our study addresses this gap by establishing the correlation between KLF7 and CSC traits in OSCC, broadening the understanding of the KLF family’s diverse functions.

Cancer stemness is strongly associated with metastasis, treatment resistance, and immune evasion. Therefore, developing therapeutic strategies that target cancer stemness could substantially enhance treatment outcomes^36^. Although we have uncovered the link between KLF7 and stemness, as a transcription factor, KLF7 is considered undruggable due to its structural and functional complexities^51^. A more viable strategy is to focus on downstream molecules that play crucial roles in the KLF7-mediated regulation of stemness and are more amenable to therapeutic targeting. Using ChIP-seq and functional assays, we identified ITGA2 as a key downstream regulator of stemness influenced by KLF7.

ITGA2, a member of the integrin family, is a cell surface protein that acts as a crucial node, transmitting signals from the ECM to the intracellular environment and triggering downstream pathways^52^. Recent evidence suggests that CSCs represent a plastic cellular state, dynamically shaped by the interaction between CSCs and their niche^53^. Various components within the tumor niche, such as cancer-associated fibroblasts and immune cells, can influence stemness through mechanisms involving cytokines or metabolites^54,55^. Considering that differentiated tumor cells can acquire stem-like properties within a CSC-supportive environment, directly eliminating CSCs may not be a practical approach. Consequently, there has been growing research interest in targeting the CSC niche or disrupting the interactions between CSCs and their surrounding environment. The ECM is a key component of the tumor niche, and ECM–tumor or ECM–immune cell interactions play a crucial role in promoting CSC properties^55,56^. Therefore, targeting the ECM and its associated signaling pathways presents a promising strategy to inhibit cancer stemness.

Our findings demonstrate that ITGA2 is enriched in cancer cells with high stemness and regulates the stemness of OSCC. Similar to our findings, ITGA2 has been shown to be enriched in glioma stem cells and co-expressed with the stem cell marker SOX2^57^. Furthermore, elevated ITGA2 expression has been observed in glioblastoma CSCs compared to their differentiated counterparts^58^. Our research further suggests that upon binding to type I collagen, ITGA2 activates several crucial pathways involved in stemness, proliferation, survival, and angiogenesis. Based on these findings, we propose that ITGA2 plays a crucial role in mediating the interactions within the CSC niche. By overexpressing ITGA2, CSCs can receive more signals that reinforce their stemness, positioning ITGA2 as a promising therapeutic target.

Additionally, as a membrane protein, ITGA2 is highly amenable to drug targeting. Several agents targeting ITGA2, including blocking antibodies and small-molecule inhibitors, are available^59,60^. Our *in vitro* and *in vivo* experiments suggest that ITGA2-specific inhibitors can suppress the stemness of OSCCs and increase their sensitivity to cytotoxic drugs. We are particularly interested in exploring whether blocking antibodies targeting ITGA2 could elicit similar effects, and further, whether ITGA2 inhibitors could enhance the efficacy of immunotherapy.

Despite the robust evidence presented in this study, several limitations exist. First, most of our experiments were conducted *in vitro*, and additional *in vivo* studies are needed to provide more comprehensive evidence. Second, this study used OSCC cell lines, and future studies employing patient-derived xenografts or tumor organoids could improve the generalizability of our findings. In conclusion, we have identified KLF7 as a regulator of CSC properties in OSCC through its enhancement of ITGA2-mediated ECM-supported stemness acquisition. Moreover, we demonstrated the feasibility of targeting ITGA2 as a potential therapeutic approach.

## Method

### Single-cell RNA-seq data processing

Single-cell transcriptome sequencing data from OSCC samples were obtained from GSE234933 and GSE82227 and analyzed using the Seurat^61^ R package (version 5.0.3). Cell barcodes were filtered to include those with 1,000–7,500 genes and < 20% mitochondrial genes. The DoubletFinder^62^ package (ver. 2.0.2) was employed to identify potential doublets. In addition, sctransform was applied for normalization, and the Harmony algorithm^63^(version 1.2.0) was used to integrate the datasets. Copy-number variation analysis was performed using the HoneyBADGER^64^ package (ver. 0.1.14).

Principal component analysis was computed based on highly variable genes. Cell clustering was conducted using the FindNeighbors and FindClusters functions, and the RunUMAP function was used for visualization when appropriate. Differentially expressed genes were identified using the FindMarkers function with the test.use parameter set to “presto.” Gene Set Variation Analysis (ver. 1.50.1)^65^ was employed to estimate pathway activities, with differences in pathway activity scores calculated using the limma package (ver. 3.58.1). Cell trajectory inference was performed using the Monocle 3 (ver. 1.3.6)^66^ packages. The CytoTRACE 2^67^ tool was used to compute potency scores and assess the state of cell differentiation. AUCell^68^ was used to measure stemness and invasiveness activity.

### TCGA analysis

RNA sequencing and corresponding clinical data for head and neck squamous cell carcinoma were downloaded from The Cancer Genome Atlas (TCGA) database (https://www.cancer.gov/ccg/research/genome-sequencing/tcga). We extracted RNA-seq data from 334 primary tumor specimens, mainly located in the oral cavity, and 44 normal specimens. Additionally, RNA-seq data and clinical information from 33 tumor types were obtained from the TCGA database using Xena (https://xenabrowser.net) for pan-cancer analysis. Wilcoxon tests were performed to analyze significant differences. Kaplan–Meier survival curves were used to assess survival differences between groups, with the log-rank test used to determine statistical significance.

### Cell culture

OSCC cell lines (CAL27 and HSC3) and HEK293T cells were obtained from the Oral Laboratory of Zhejiang University. All cell lines were cultured in Dulbecco’s modified Eagle medium (DMEM) supplemented with 10% fetal bovine serum and 1% penicillin/streptomycin and maintained at 37°C in 5% CO2. Cells were passaged at 70–90% confluency, and mycoplasma contamination was regularly checked using a detection kit.

#### Samples and animals

Formalin-fixed, paraffin-embedded patient samples were retrospectively obtained from the First Affiliated Hospital of Zhejiang University. NOD/SCID nude rats were purchased from the Second Affiliated Hospital of Zhejiang University. The study adhered to the Declaration of Helsinki, and the study protocol was approved by the Ethics Committee of both the First and Second Affiliated Hospitals of Zhejiang University.

### Plasmids and lentiviral transfection

The KLF7 overexpression and knockdown lentiviral vectors were obtained from Vigene (Shandong, China), and the ITGA2 lentiviral vector was sourced from GenePharma (Shanghai, China). The promoters of wild-type and mutant ITGA2 were cloned into the pGL4.19 vector. The ITGA2 plasmid was synthesized in our laboratory: full-length human ITGA2 was amplified from HEK293T cDNA and cloned into the Xho I and EcoR I restriction sites of the pcDNA3.1-Flag vector using the ClonExpress MultiS One Step Cloning Kit (#C113-02; Vazyme). ITGA2 mutants were generated by site-directed mutagenesis using the Mut Express MultiS Fast Mutagenesis Kit (#C215-01; Vazyme) according to the manufacturer’s instructions. To establish lentiviral transfected cell lines, lentiviruses were added to CAL27 and HSC3 cells, and 24 h later, the medium was replaced with medium containing 2 µg/mL puromycin. Plasmid transfections were carried out using Lipofectamine 3000 (Invitrogen, Carlsbad, CA, USA) according to the manufacturer’s instructions. The primer sequences are presented in the Supplementary Tab.3.

### Scratch wound assay

For scratch wound assays, cells were seeded into 6-well plates and grown to confluence. A 200-µL pipette tip was used to create a scratch wound, followed by two washes with phosphate-buffered saline to remove detached cells. Serum-free medium was added, and wound closure was monitored under the microscope every 3 h. ImageJ software was used to analyze wound closure.

### Transwell migration assay

Serum-free medium was used to prepare HSC3 and CAL27 cell suspension, 200µL of cell suspension was added to the upper chamber of the Transwell (3403, Corning) while 800 µL of complete culture medium was added to the lower chamber. After incubation for 48 hours, 4% paraformaldehyde-fixed the cells that passed through the cell chamber filter attached to the surface and stained with 0.4% crystal violet.

### Immunohistochemistry and immunofluorescence

All paraffin-embedded tissues were sectioned into 4-μm slices. After deparaffinization, rehydration, antigen retrieval, and blocking, the sections were incubated overnight with primary antibodies. After incubation with secondary antibodies, sections were stained with 3,3′-diaminobenzidine and counterstained with hematoxylin. For tissue immunofluorescence, after routine blocking and incubation with primary and secondary antibodies, tyramide signal amplification staining was performed. Subsequently, second and third antibodies were incubated, followed by DAPI staining. The antibodies used in this study included KLF7(ab197690,Abcam), ITGA2 (A19068,ABclonal),and CD44 (A21919, Abclonal). For cellular immunofluorescence, cells were plated on coverslips and fixed with paraformaldehyde upon reaching 30% confluence, followed by permeabilization. After washing and blocking, the primary antibody against YAP1 (66900-1-Ig, Proteintech) was incubated overnight. The slides were then incubated with the anti-mouse secondary antibody(ab150115, Abcam) and counterstained with DAPI.

### Immunoblotting

Cells were lysed in RIPA buffer containing protease inhibitors. After scraping, lysates were centrifuged, and supernatants were collected. Protein concentrations were measured using a BCA kit. After Sodium dodecyl sulfate-polyacrylamide gel electrophoresis (SDS-PAGE) and membrane transfer, blocking was performed, and primary antibodies were incubated overnight at 4°C. The primary antibodies used in this study included anti-KLF7(ab197690, Abcam), anti-ITGA2(A19068, ABclonal), anti-E-cadherin(ab40772, Abcam), anti-vimentin (A19607, Abclonal), anti-β-actin (8457S, CST), anti-NANOG(D73G4, CST), anti-AKT(9272S, CST), anti-p-AKT(4060S, CST), anti-ERK1/2(4695S, CST), anti-p-ERK1/2(4370S, CST), anti-YAP1(66900-1-Ig, Proteintech), anti-p-YAP1(29018-1-AP, Proteintech), and anti-COL1A1 (A25310,Abclonal).

### Immunoprecipitation

For immunoprecipitation, cells or tissues were homogenized in lysis buffer. Protein extracts were incubated with Pierce™ anti-DYKDDDDK Magnetic Agarose (Thermo Fisher Scientific). The binding complexes were washed with buffer, mixed with loading buffer, and subjected to SDS-PAGE. The presence of ITGA2 in the precipitates was detected using a rabbit anti-ITGA2 antibody(A19068, ABclonal), and collagen 1 was detected using an anti-COL1A1 antibody (A25310, Abclonal).

### Sphere formation assay

To assess sphere formation, cells were collected and suspended in serum-free DMEM/F12 supplemented with 20 ng/mL epidermal growth factor(PMG8045, Thermo Scientific), 20 ng/mL basic fibroblast growth factor(13256029, Thermo Scientific), and 1× B27(17504044, Thermo Scientific). The single-cell suspension was seeded at 2,000 cells per well in suspension culture and maintained for 7–10 days. Sphere sizes were observed under a microscope.

### Flow cytometry

For flow cytometry, cells were collected and resuspended at a concentration of 1 × 10^6^ cells per 200 µL of phosphate-buffered saline per tube. Cells were incubated with CD133 antibody (566596, BD Pharmingen) on ice for 30 min, washed two or three times, and analyzed.

### Limiting dilution assay

In limiting dilution assays, cells were suspended at various densities (1 × 10^6^, 5 × 10^5^, 1 × 10^5^, and 5 × 10^4^ cells per 100 µL) in serum-free DMEM containing 10% Matrigel and injected into 6-8-week-old female nude mice. Tumor growth was monitored, and after 3-4 weeks, the mice were sacrificed for sample collection.

### ChIP-seq analysis

The establishment of MYC-KLF7+-expressing CAL27 cells and ChIP-seq were performed by Igenebook Biotechnology Co. Ltd. (Wuhan, China). Briefly, 1 × 10^7^ cells were washed and cross-linked with 1% formaldehyde, and then quenched by adding glycine. Cells were lysed, and chromatin was isolated on ice. Chromatin was sonicated to produce soluble, sheared chromatin. Next, 20 µL of chromatin was saved for input DNA, whereas 100 µL of chromatin was used for immunoprecipitation with anti-MYC antibodies (D3N8F, CST). The immunoprecipitated material was bound to protein beads, washed, and eluted from the beads. The eluted material was treated with RNase A followed by proteinase K. The immunoprecipitated DNA was used to construct sequencing libraries, which were sequenced on an NovaSeq 6000 (Illumina, San Diego, CA, USA) using paired-end 150 bp sequencing.

### ChIP-seq data analysis

Trimmomatic (ver. 0.36) was used to filter out low-quality reads^69^. Clean reads were mapped to the human genome using BWA (ver. 0.7.15)^70^. Samtools (ver. 1.3.1) was applied to remove potential PCR duplicates ^71^. Peak calling was performed using MACS2 (ver. 2.1.1.20160309) with default parameters (bandwidth, 300 bp; model fold, 5, 50; q value, 0.05). Peaks were assigned to genes if their midpoints were closest to the transcription start site of a gene^72^. Motif occurrence within peaks was predicted using HOMER (ver. 3) with a maximum motif length of 12 base pairs^73^. The clusterProfiler^74^ (http://www.bioconductor.org/packages/release/bioc/html/clusterProfiler.html) package in R software [was employed to perform Gene Ontology (http://geneontology.org/)^75^ and Kyoto Encyclopedia of Genes and Genomes (http://www.genome.jp/kegg/) enrichment analysis^76^, with a hypergeometric distribution and a q value cutoff of 0.05^74^.

### Luciferase reporter assays

For the luciferase reporter assays, CAL27 cells were seeded onto 24-well plates and transfected with pGP4.19-ITGA2 promoter, mutant constructs, and pRL-TK vector, when co-transfected with KLF7+ vector or control for 24 h. Cells were then harvested and lysed using lysis buffer (Promega Biosciences) at room temperature for 15 min. Luciferase activity was measured using the Dual Luciferase Reporter Assay System Kit (Promega Biosciences) according to the manufacturer’s instructions. Total light production (optical density: 490 nm) was measured using the SpectraMax M3 multimode microplate reader (Molecular Devices) and normalized to Renilla luciferase activity.

### Xenograft

For *in vivo* experiments, tumor cells were suspended in serum-free DMEM and injected subcutaneously into the axillae of 4–6-week-old nude mice. Tumor growth was measured using calipers every 3 days and recorded in cubic centimeters. In inhibitor treatment experiments, cisplatin (A821, APExBIO, 5 mg/kg) or TC-I 15 (HY-107588, MCE, 20 mg/kg) was injected intravenously on days 6, 9, 12, 15, and 18 post-cell injection. Mice were euthanized when the tumor size exceeded a predetermined threshold for survival analysis. All mice were housed under pathogen-free conditions at Zhejiang Chinese Medical University, and the study protocol was approved by the university’s ethics committee.

### Data analysis

Pooled data are presented as the mean ± standard error of mean, unless otherwise indicated. Detailed information on sample sizes, error bars, and statistical analyses can be found in the figure legends. P values for statistical comparisons between two groups or multiple groups were calculated using Excel 2021 (Microsoft) and GraphPad Prism (ver. 9.2.0) (GraphPad Software).

## Acknowledgements

This work was supported by the National Natural Science Foundation of China [81902760]

## Contributions

Jiong Lyu and Xin Qi designed the research and Jiong Lyu, Huiyong Zhu, and Cang Li supervised the study. Xin Qi, Jiang Zhou, and Pan Wang designed and performed the initial experiments. Xin Qi, Wang Pan and Yunyan Li performed the data analysis. Xin Qi, Haoran Li, Yuwen Miao, XiaoQing MA, Yanling He, and Zhiling Zhang performed most of the biochemical experiments. Xin Qi performed animal experiments. Xin Qi and Jiong Lyu wrote the manuscript with contributions from Rutao Cui, Wenquan Quan, Xiayan Luo, and Wenyi Shen.

