## Supplementary Fig.1 for "KLF7-Regulated ITGA2 as a Therapeutic Target for Inhibiting Oral Cancer Stem Cells"

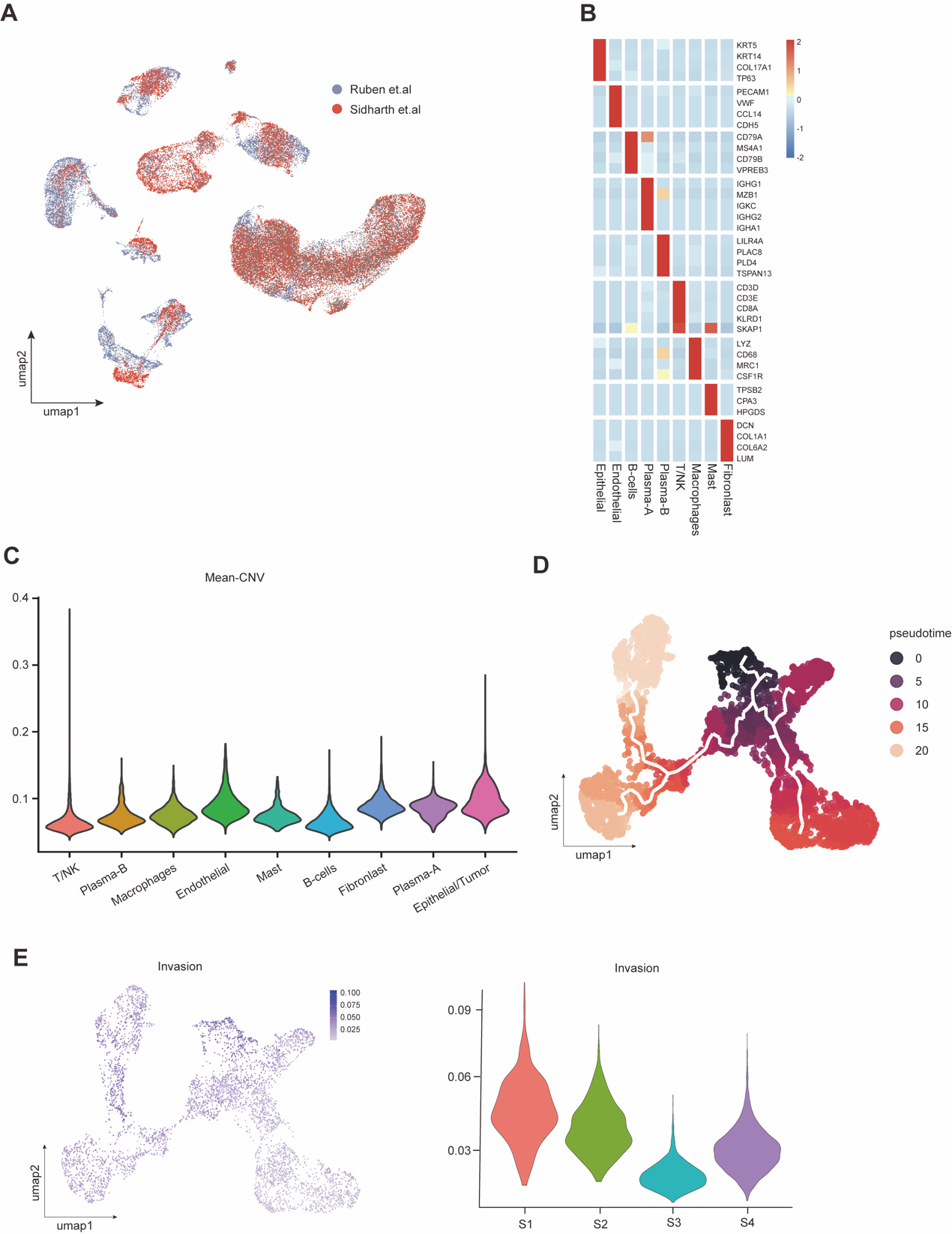

**Supplementary Fig.1: Related to Fig.1**

**A** UMAP plot of all cells, colored by OSCC sample source. **B** Heat map showing expression of selected marker genes by all cells assigned to each cell type. **C** Violin plot depicting mean Copy Number Variation (CNV) scores for the different cell types. **D** Pesuedotime plot of the subtype of malignant cells. **E** Invasive gene sets activity (AUCell score) in all malignant cells (umap plots, left panel) or per subtype of malignant cells (violin plots, right panel)
