## Supplementary Fig.2 for "KLF7-Regulated ITGA2 as a Therapeutic Target for Inhibiting Oral Cancer Stem Cells"

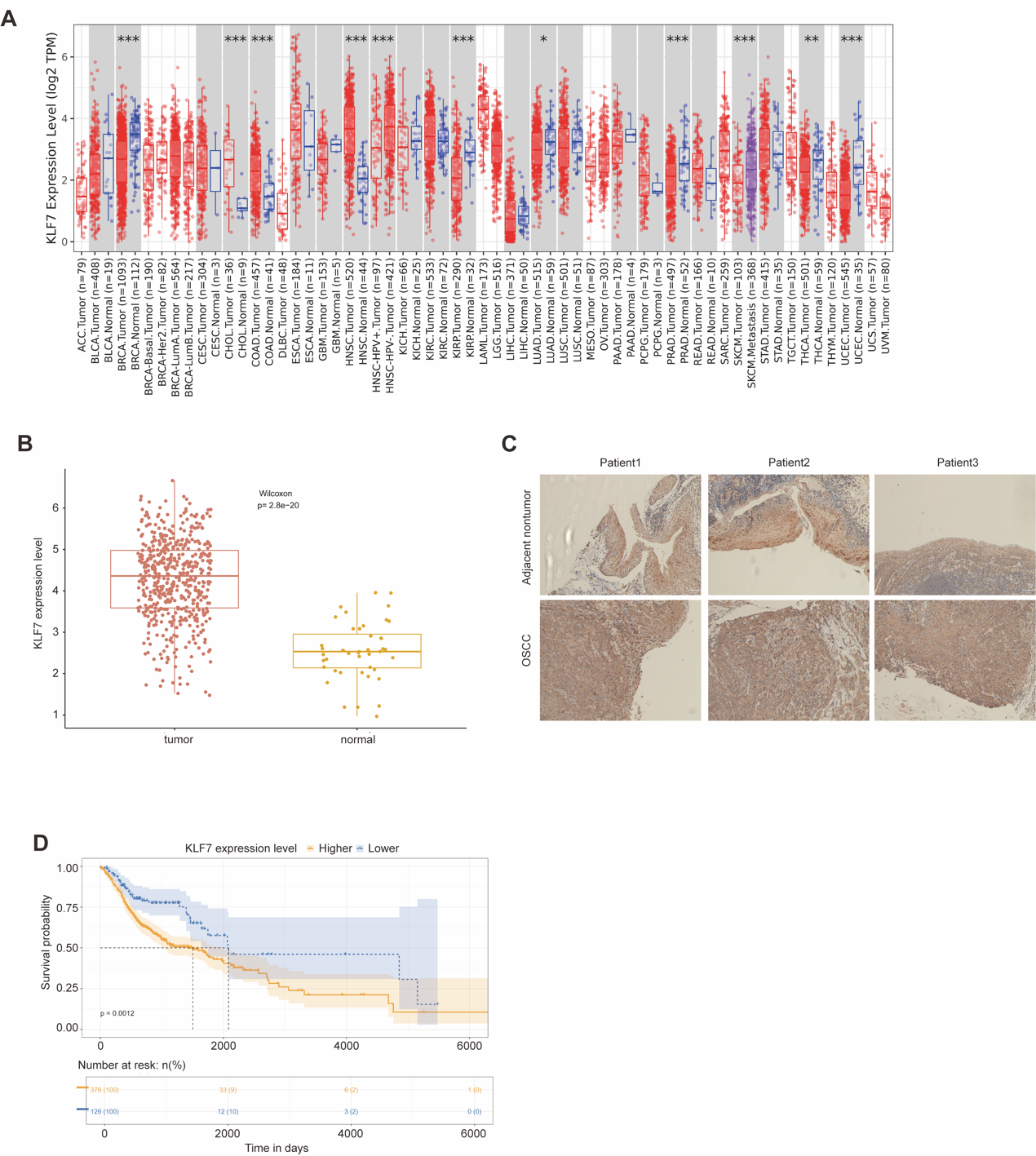

**Supplementary Fig.2: KLF7 overexpress in OSCC and related to poor prognosis**

**A** KLF7 expression in pan-cancer in TCGA dataset. **B** KLF7 expression in OSCC and normal samples. **C** Representative immunohistochemistry image showing the expression of KLF7 in human OSCC sample. **D** Kaplan-Meier curve of KLF7 in OSCC TCGA dataset
