## Supplementary Fig.3 for "KLF7-Regulated ITGA2 as a Therapeutic Target for Inhibiting Oral Cancer Stem Cells"

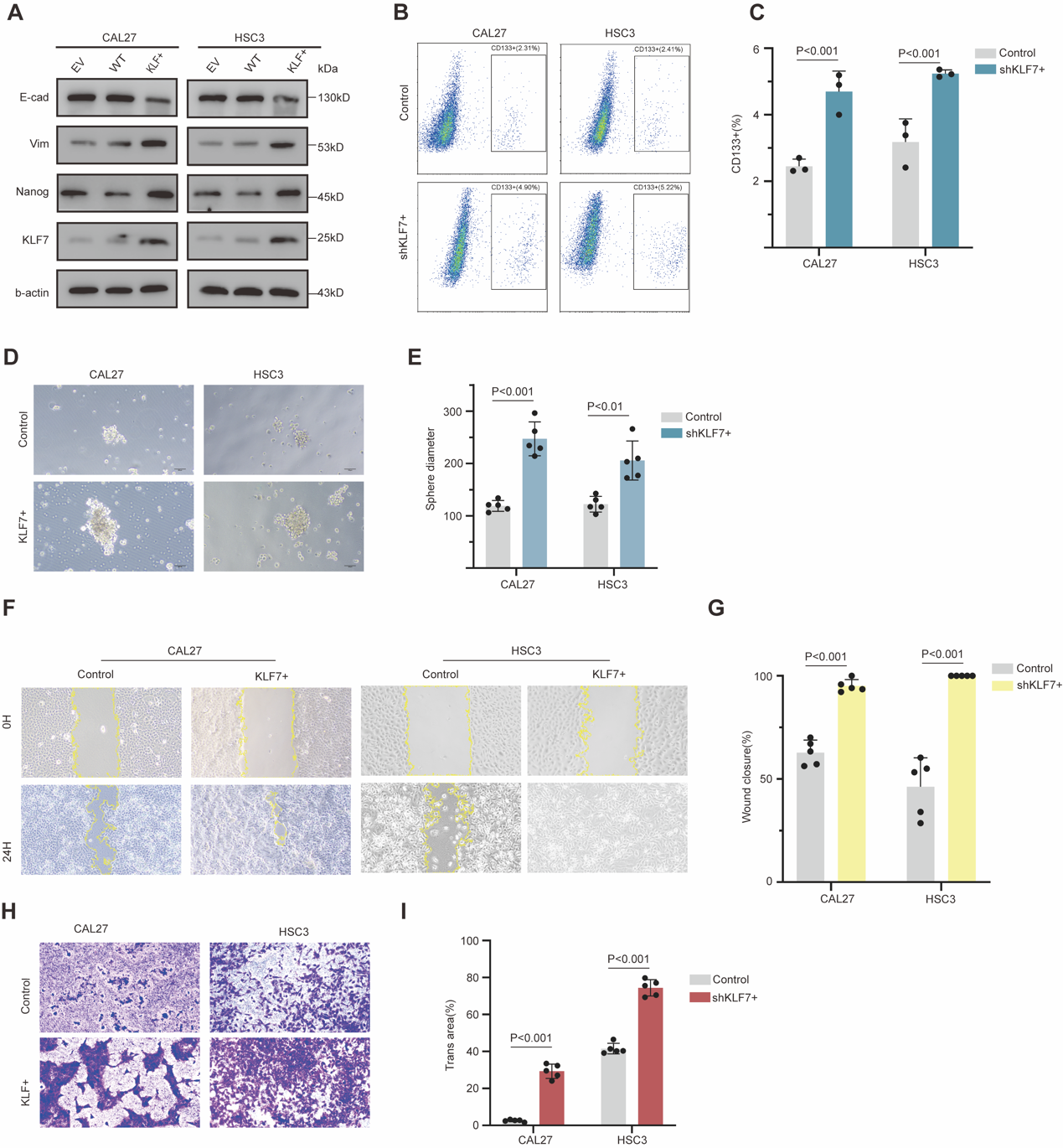

**Supplementary Fig.3: Overexpress KLF7 increases the stemness and metastasis of OSCC**

**A** Immunoblot assay of KLF7, NANOG, VIM, and e-cad protein levels in CAL27 and HSC3 cells after stable overexpressing KLF7 in CAL27 and HSC3 cells. **B** Identification of OSCC CSCs (CD133+) in wt and overexpress KLF7 cells by flow cytometry (n = 3). **C** Quantification of FACS analysis in B. **D** Representative image of Sphere formation assay in wt and overexpress KLF7 cells in normal sphere medium (n = 5). **E** Quantification of spheres diameter in D. **F** Representative image of wound-healing image after 24h in wt and overexpress KLF7 cells (n=5). **G** Quantification of percentage of wound closure in F. **H** Representative image of transwell after 24h in wt and overexpress KLF7 cells (n=5). **I** Quantify the percentage of trans area in H.
