## Supplementary Fig.4 for "KLF7-Regulated ITGA2 as a Therapeutic Target for Inhibiting Oral Cancer Stem Cells"

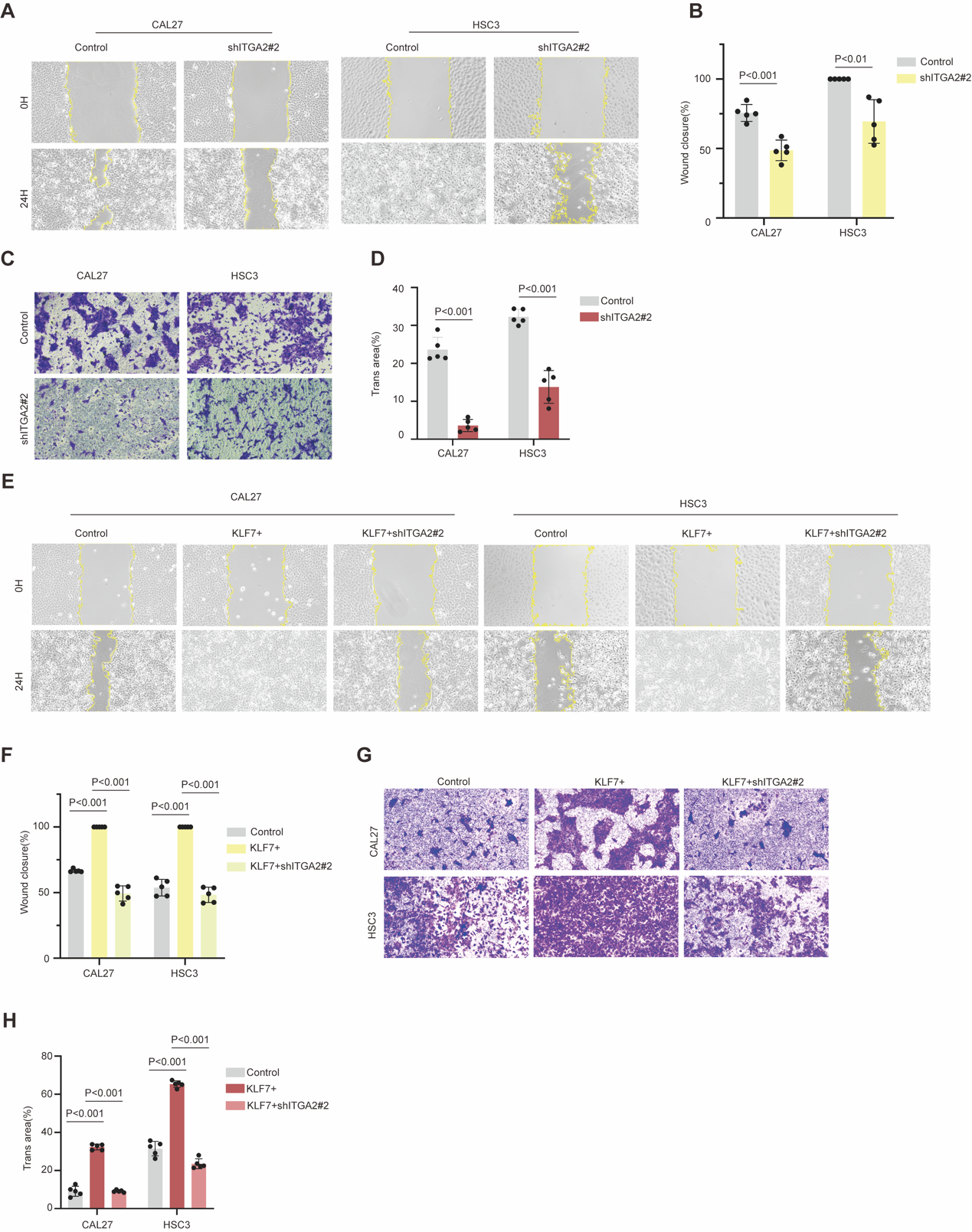

**Supplementary Fig.4: Knockdown ITGA2 inhibits the metastasis of OSCC**

**A** Representative image of wound-healing image after 24h in wt and silencing ITGA2 cells (n=5). **B** Quantification of the percentage of wound closure in A. **C** Representative image of transwell after 24h in wt and silencing ITGA2 cells (n=5). **D** Quantify the percentage of trans area in C. **E** Representative image of wound-healing image after 24h in wt, overexpress KLF7 and overexpress KLF7 followed by knockdown ITGA2 cells (n=5). **F** Quantification of percentage of wound closure in E. **G** Representative image of transwell after 24h in wt, overexpress KLF7 and overexpress KLF7 followed by knockdown ITGA2 cells (n=5). **H** Quantify the percentage of trans area in G.
